# Dynamics of a neuronal pacemaker in the weakly electric fish *Apteronotus*

**DOI:** 10.1101/2020.06.26.173088

**Authors:** Aaron R. Shifman, Yiren Sun, Chloé M. Benoit, John E. Lewis

## Abstract

The precise timing of neuronal activity is critical for normal brain function. In weakly electric fish, the medullary pacemaker network (PN) sets the timing for an oscillating electric organ discharge (EOD) used for electric sensing. This network is the most precise biological oscillator known, with sub-microsecond variation in oscillator period. The PN consists of two principle sets of neurons, pacemaker and relay cells, that are connected by gap junctions and normally fire in synchrony, one-to-one with each EOD cycle. However, the degree of gap junctional connectivity between these cells appears insufficient to provide the population averaging required for the observed temporal precision of the EOD. This has led to the hypothesis that individual cells themselves fire with high precision, but little is known about the oscillatory dynamics of these pacemaker cells. To this end, we have developed a biophysical model of a pacemaker neuron action potential based on experimental recordings. We validated the model by comparing the changes in oscillatory dynamics produced by different experimental manipulations. Our results suggest that a relatively simple model captures the complex dynamics exhibited by pacemaker cells, and that these dynamics may enhance network synchrony and precision.

**Author summary:** Many neural networks in the brain exhibit activity patterns which oscillate regularly in time. These oscillations, like a clock, can provide a precise sense of time, enabling drummers to maintain complex beat patterns and pets to anticipate “feeding time”. The exact mechanisms by which brain networks give rise to these biological clocks are not clear. The pacemaker network of weakly electric fish has the highest precision of all known biological clocks. In this study, we develop a detailed biophysical model of neurons in the pacemaker network. We then validate the model against experiments using a nonlinear dynamics approach. Our results show that pacemaker precision is due, at least in part, to how individual pacemaker cells generate their activity. This supports the idea that temporal precision in this network is not solely an emergent property of the network but also relies on the dynamics of individual neurons.

## Introduction

Timing of neuronal spikes is critical to many brain processes, including sound localization [1–3], escape responses [4–6], and learning and memory [7,8]. When neural processes are periodic, they can form the basis for biological clocks which span a range of precision (variability in oscillation period), with a higher variability leading to a less reliable clock.

Variability in the period of neuronal oscillators (reported as a coefficient of variation: CV = s.d./mean) can be relatively high, as in the bullfrog sciatic nerve with a CV=0.37 [9,10]. For reference, a random Poisson process has a CV=1, while neurons in the visual system can have a CV >1 [11,12]. In contrast, the neural oscillators underlying the electric organ discharge (EOD) of the weakly electric fish *Apteronotus* have a CV as low as ~10^−4^ (corresponding to a raw standard deviation of ~100ns), making it the most precise biological oscillator known [10,13]. The high precision of the EOD of *Apteronotus* makes it a particularly attractive model for the study of neural circuit timing [10,13,14].

*Apteronotus* generates an oscillating electric field (EOD) to sense their environment in the dark [15]. Heterogeneities in the environment perturb the EOD, and these perturbations are sensed by electroreceptors on the skin. The timing of the oscillations underlying the EOD are set by the medullary pacemaker network (PN) [13, 16–19]. This nucleus is a collection of two principle cell types: pacemaker cells which are intrinsic to the PN, and relay cells which project down the spinal cord to drive the EOD [14,16,20]. Additionally, there are parvalbumin positive cells (parvocells) whose function is currently unknown, but are not thought to contribute to the oscillatory function of the PN [21].

The pacemaker cells in the PN are highly synchronized, with relative phases across cells close to 2% of the oscillator period [10,14]. In general, networks are thought to achieve high-precision and high-synchrony through the population-averaged activity of a large number of strongly-connected cells [14,16,22]. However, pacemaker and relay cells are connected only sparsely, with weak gap junctions [14, 16–18]. Although network connectivity may be functionally enhanced through the electric feedback from the EOD itself [14,23], the apparent disconnect between high-synchrony or high-precision and low connectivity in the PN may also be explained by the high precision of individual cells [16], with synchrony emerging from weak interactions between precise cells with stereotyped dynamics. Indeed, some underlying oscillatory dynamics are thought to be more amenable to synchronization than others [24–26].

Previous studies have used a Hodgkin-Huxley based model to explore PN synchrony and precision [14,16], but this model was not intended to accurately represent the action potential waveform of pacemaker neurons. While these studies provided insight into pacemaker network interactions, a more accurate biophysical model is required to determine how transmembrane currents, intrinsic oscillatory dynamics and gap junctional coupling impact single cell precision and network synchrony. To this end, we present a biophysically based pacemaker cell model which accurately captures the waveform of pacemaker cells as well as their dynamical responses to experimental manipulations.

## Methods

### Model Developement

Previous pharmacological experiments have suggested that pacemaker cells express the following suite of ionic currents: inactivating sodium (I_Na_) and/or persistent sodium (I_Na_P__), inactivating potassium (I_K_), T/R type calcium (I_Ca_), and leak (I_L_) [27]. Using the standard Hodgkin-Huxley-style biophysical approach [28], these currents underlie the dynamics of our model (see supplemental equations [S1 File] for details). Model parameters were fit to intracellular recordings from previously published experiments on *Apteronotus leptorhynchus* pacemaker cells [14]. A standard waveform from a representative pacemaker cell comprising two successive action potentials, averaged over 30 sweeps, was used to fit the primary model. Fitting two successive action potentials (oscillator cycles), rather than one, minimizes frequency drift between model and target waveforms. We also show that the primary model can be generalized by fitting it to waveforms from other pacemaker cells in both *Apteronotus leptorhynchus* and the related species *Apteronotus albifrons* (see figure 1). Note that Smith and Zakon (2000) also suggested a role for a persistent sodium current (I_Na_p__), but our initial studies showed that including this current resulted in many solutions that would not spike (not shown), so we did not include I_Na_p__ in our final model. This current may provide a means of modulating frequency but it is not necessary to explain variations in pacemaker cell waveform across individuals and species.

**Fig 1.**
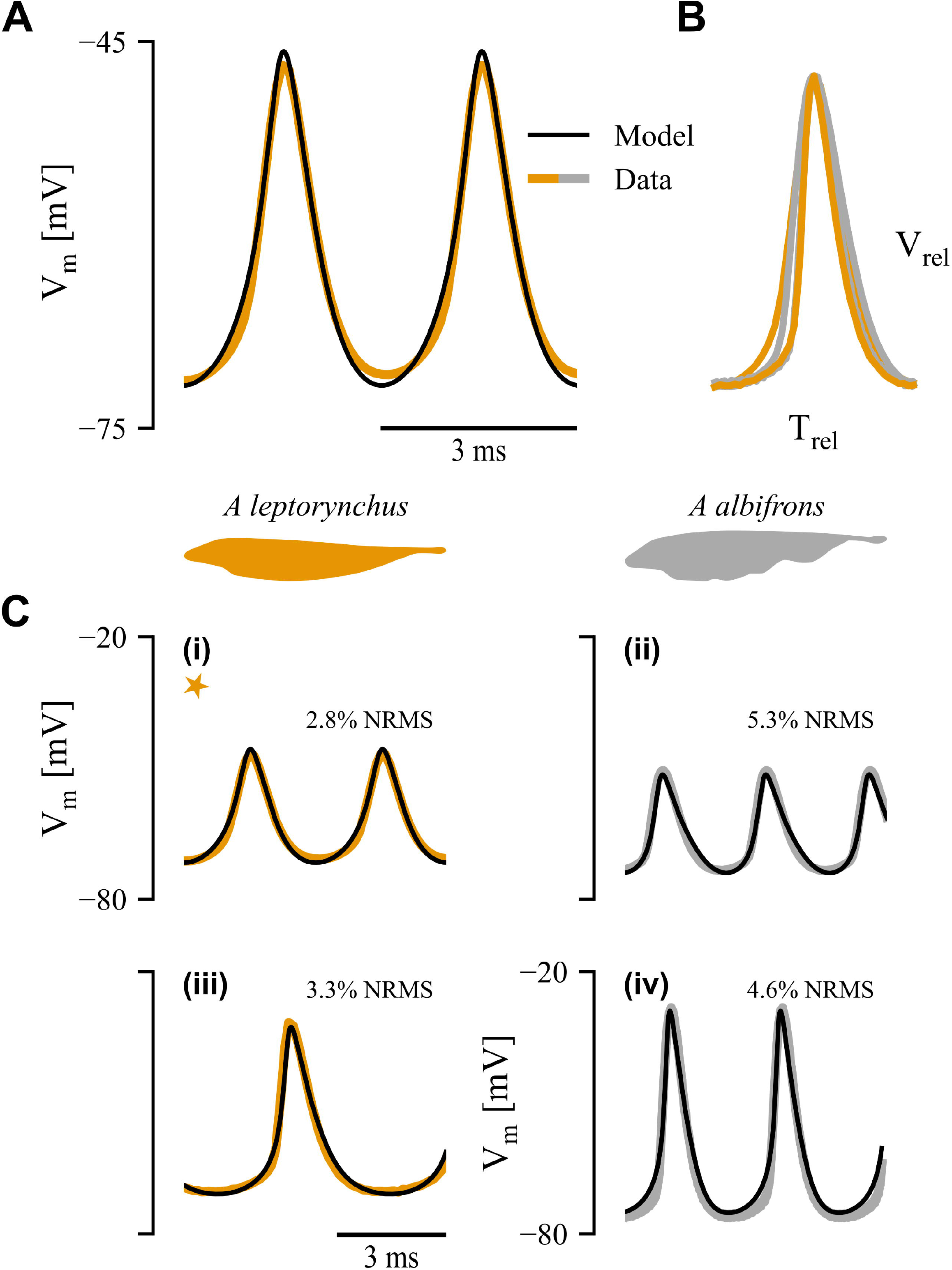
Model fit results. (A) Canonical model fit (black line) to *A. leptorhynchus* action potential waveform (orange). (B) Dimensionless waveform (action potential normalized by period in time and peak-peak amplitude) from two individuals from each species. (C) Data fits showing model flexibility over a range of frequencies, amplitudes and means for *A. leptorhynchus* (left, orange) and *A. albifrons* (right, grey). Orange star indicates model fit in panel A.

We implemented a generic, parameterized model in the Brian2 simulation engine (version 2.3) [29], and fit our model using a differential evolutionary algorithm provided in the brian2modelfitting package (version 0.3). This algorithm is similar to a genetic algorithm and starts with a large set of parameters drawn randomly within set bounds. Based on the fitting performance (i.e. “fitness”; see later discussion on fitting error), some parameter values will have a higher or lower probability of being used in the next iteration. Stochastic perturbations within the parameters allow for an efficient sampling of large parameter spaces [30,31]. The algorithm was initialized with 5000 samples of each parameter and run for 3 iterations. Each parameter was sampled uniformly between upper and lower bounds, based roughly on known biophysical principles (see S1 Table).

Fitting error was quantified using the root mean squared (RMS eq. 1) error when the two waveforms (experimentally measured V^e^, and model V^m^) were aligned by first spike times (defined as the action potential peak), 〈*x*〉 represents the mean of *x*.

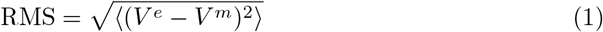

### Experimental methods

To validate the model, brain slices of the pacemaker nucleus were prepared as described previously [10,14]. Briefly, adult black ghost knifefish (*A. albifrons*) were obtained from commercial fish suppliers and housed on a 12/12 light-dark cycle in flow-through tanks at water temperature 27-28°C and conductivity 150-250μS. All housing and experimental protocols were in accordance with guidelines approved by the Animal Care Committee of the University of Ottawa (BL-1773). Fish (N=5) were deeply anaesthetized using 0.1% Tricaine methanosulfate (TMS, Syndel International Inc, Nanaimo, BC, Canada) before being transferred to a bath of ice-cold artificial cerebrospinal fluid (ACSF; in mM: 124 NaCl, 24 NaHCO_3_, 10 D-Glucose, 1.25 KH_2_PO_4_, 2 KCl, 2.5 MgSO_4_, 2.5 CaCl_2_; bubbled with 95% O_2_/5% CO_2_). The brain was quickly removed and the pacemaker nucleus cut away using fine scissors (~1mm rostral, 2mm caudal, 1mm dorsal) and transferred to a 35mm petri dish perfused with oxygenated room-temperature (22 °C) ACSF. After a minimum of 30 minutes, pacemaker recordings (intracellular or extracellular) were performed with borosilicate glass sharp electrodes (30-90MΩ, P-2000 electrode puller, Sutter Instrument Company, Novato, CA, USA) using an Axoclamp 2B amplifier (Molecular Devices, Sunnyvale, CA, USA). Data was acquired using a Digidata 1440a digitizer (Molecular Devices) at a sampling frequency of 100kHz using pClamp 10 (Molecular Devices). Low Na^+^ ACSF was prepared in a similar fashion as ACSF, only substituting NaCl for equimolar amounts of sucrose (Fisher Chemical, Fair Lawn, NJ, USA). The perfusion system involved a transfer time of approximately 4 minutes when switching between Na^+^ and low-Na^+^ ACSF solutions.

Action potential frequency in the PN was measured from one-second recordings taken at 20s intervals using Fourier analysis (as the highest power, dominant frequency). Cessation of spiking was determined when the power at the dominant frequency was less than 1.5 times the power at 60 Hz (signal-to-noise ratio, SNR<1.5) This criterion was additionally used to identify overly noisy recordings. We additionally identify noisy recordings by ensuring that the dominant frequency is not within 5Hz of a power line harmonic.

## Results

### Model Fit

We developed a biophysical Hodgkin-Huxley-based model of a pacemaker neuron in the PN of a weakly electric fish. Motivated by previous studies [27], our model included voltage-dependent sodium, potassium, and calcium channels, along with leak channels (I_Na_, I_K_, I_ca_, I_L_). We used a differential evolutionary algorithm to survey a 44-dimensional parameter space (see Methods and Appendix). After optimization, the RMS error between model and data waveform was 0.7mV; when normalized by action potential amplitude, this corresponds to a 2.8% error. Note that small differences in action potential timing can lead to relatively large errors due to the fast rise and fall times that are typical of action potentials, so the model matches the data even better than the RMS error would suggest over most of the action potential cycle (figure 1A). For model fits see S2 Table and S3 Table.

At this point, our model describes an action potential of a single cell from an individual *A. leptorhynchus*. It is also of interest to determine how well this model will generalize across individuals and the related species *A. albifrons*. In figure 1B, we show action potential waveforms (dimensionless, normalized in both time and amplitude) from pacemaker cells from two individuals of each species (figure 1B); the similarity across waveforms suggests that the underlying dynamics are also similar. To demonstrate this and to show the flexibility of the model, we refit the model to each of these four action potential waveforms with the same parameter bounds (figure 1C; data from figure 1A is indicated by gold star). Over a range of amplitudes and frequencies, the model fits involved a worst-case error of 5.3%. And importantly, there were no systematic differences in the voltage dependence of the gating variables across all models (S1 Fig).

Previous results suggest that while calcium can contribute to action potential waveform shape, it does not fundamentally underlie pacemaker cell oscillation [27]. We tested this in our model by setting g_Ca_ to 0 (I_ca_-Block) and found that changes to the waveform were subtle and the model continues to oscillate (figure 2A). In figure 2B, we show the contributions of each current to the action potential waveform. As expected, the depolarization of the action potential is driven by the sodium current, and the repolarization/hyperpolarization is driven by potassium current, but I_Ca_ has little effect (compare the full model with the I_Ca_-Block model, Figure 2B). Overall, our modeling results confirm previous experimental results suggesting a minimal role of calcium in the pacemaker action potential oscillation.

**Fig 2.**
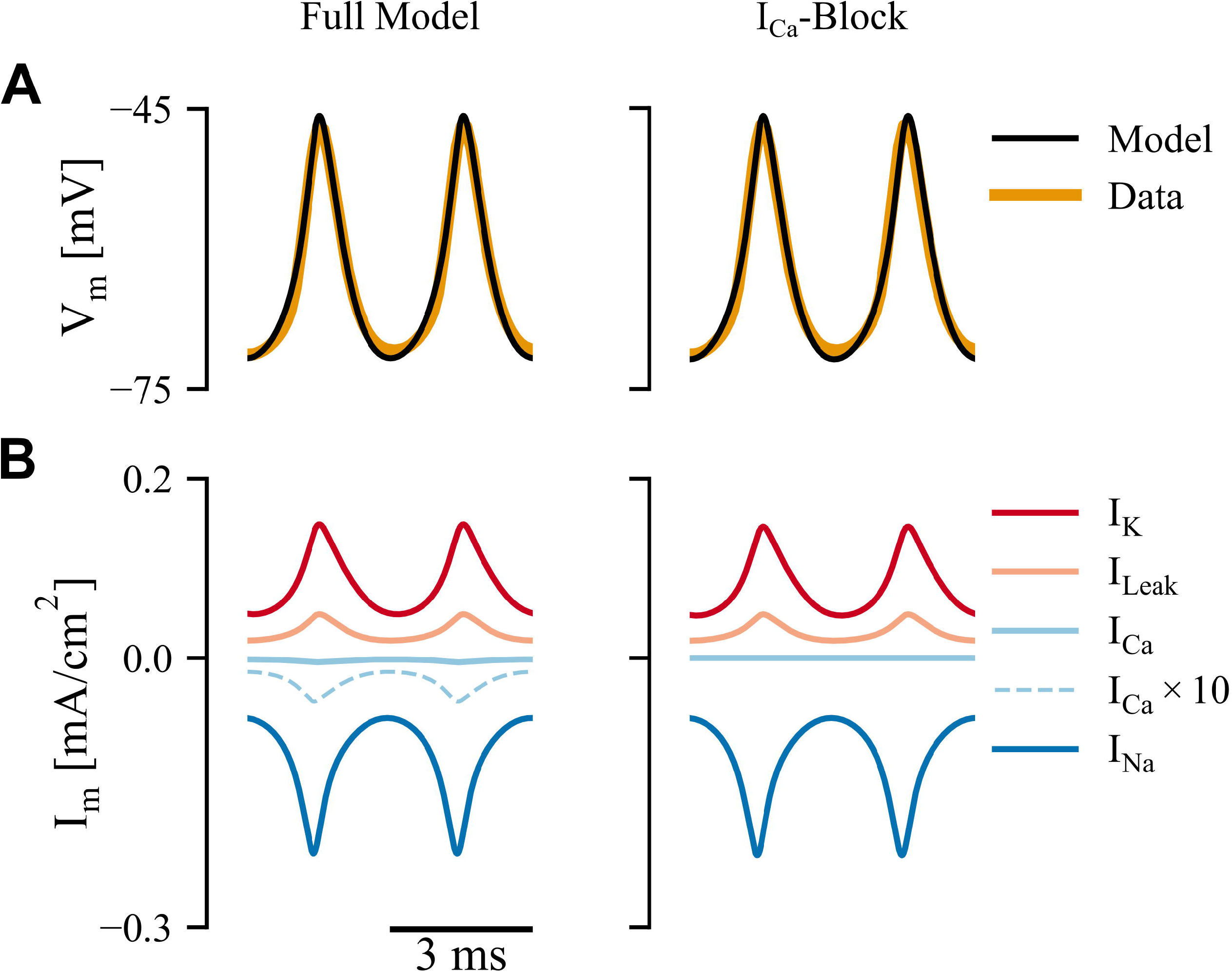
Analysis of membrane currents in the canonical model and the effects of I_Ca_-block. (A) Model fits for both full model (left) and with I_Ca_ blocked (right) showing no systematic differences. (B) Current breakdown with I_Ca_×10 (dashed light blue line) showing a 10× magnified calcium current for illustrative purposes.

At this stage, it would not be inappropriate to consider the model overfit. We have used a biophysical model with 44 parameters to fit a single oscillator waveform. A good fit would not be surprising, so it is important to validate the dynamics of our model against additional data. To this end, we consider the pacemaker dynamics during two different experimental manipulations: low concentration of extracellular sodium (decreased E_Na_), and pharmacological block of Na^+^ and K^+^ channels [21].

### Model Validation: Effects of E_Na_

To test the new pacemaker model, we compared its dynamics under conditions that differed from those in the model fitting process. In the study of dynamical systems, qualitative changes in behavior produced by small changes in a system parameter are referred to as bifurcations [32]. One particular example of a bifurcation relevant to pacemaker dynamics is the transition between an oscillating state and a rest state (non-oscillating), and vice-versa; the nature of this transition depends on the system properties as well as the particular parameter that is varied.

One classic way in which oscillations can arise is through a Hopf bifurcation. The hallmark of a Hopf bifurcation is that the transition from an oscillating state to rest (or vice-versa) involves a discontinuous jump (i.e. as a system parameter is varied, there is an abrupt change in frequency from some minimum value to zero) [32] There are two kinds of Hopf bifurcations: one that exhibits a hysteresis and one that does not. Hysteresis can manifest in many ways. In the case of a Hopf bifurcation, hysteresis appears as a bistable system i.e. at a given parameter value the system can be oscillatory or not. Bifurcations presenting with hysteresis are known as *subcritical* Hopf bifurcations, whereas those without are *supercritical*. An alternative type of bifurcation (the homoclinic bifurcation) involves a continuous transition from rest to oscillating state along with a gradual change in oscillation frequency (i.e. rest can be thought of as an oscillation with infinite period) [32]. In summary, characterizing the transition between oscillating and non-oscillating states can provide a test of system dynamics [33].

Lowering the equilibrium potential of sodium (E_Na_) via changes in extracellular Na^+^ concentration typically leads to cessation of action potential generation. We thus used this parameter to explore the transition between oscillating and non-oscillating states in both our pacemaker model and in experimental pacemaker preparations. In our model, we can manipulate E_Na_ directly. In our experiments, low-Na^+^ ACSF was washed in to dilute the control ACSF, thereby gradually decreasing E_Na_. Under ideal mixing conditions, the sodium concentration of the bath should obey an exponential diffusion equation (eq 2) where *r* is the flow rate:

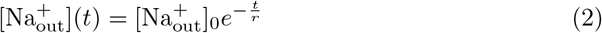

E_Na_ is given by the Nernst equation (eq 3) so by substitution we have:

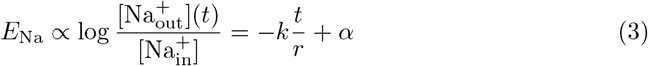

where ∝ represents proportionality, and *k* and *α* are lumped constants. This implies that E_Na_ should decrease linearly in time, and since we do not have a direct measure of E_Na_, time should be a good proxy.

**Fig 3.**
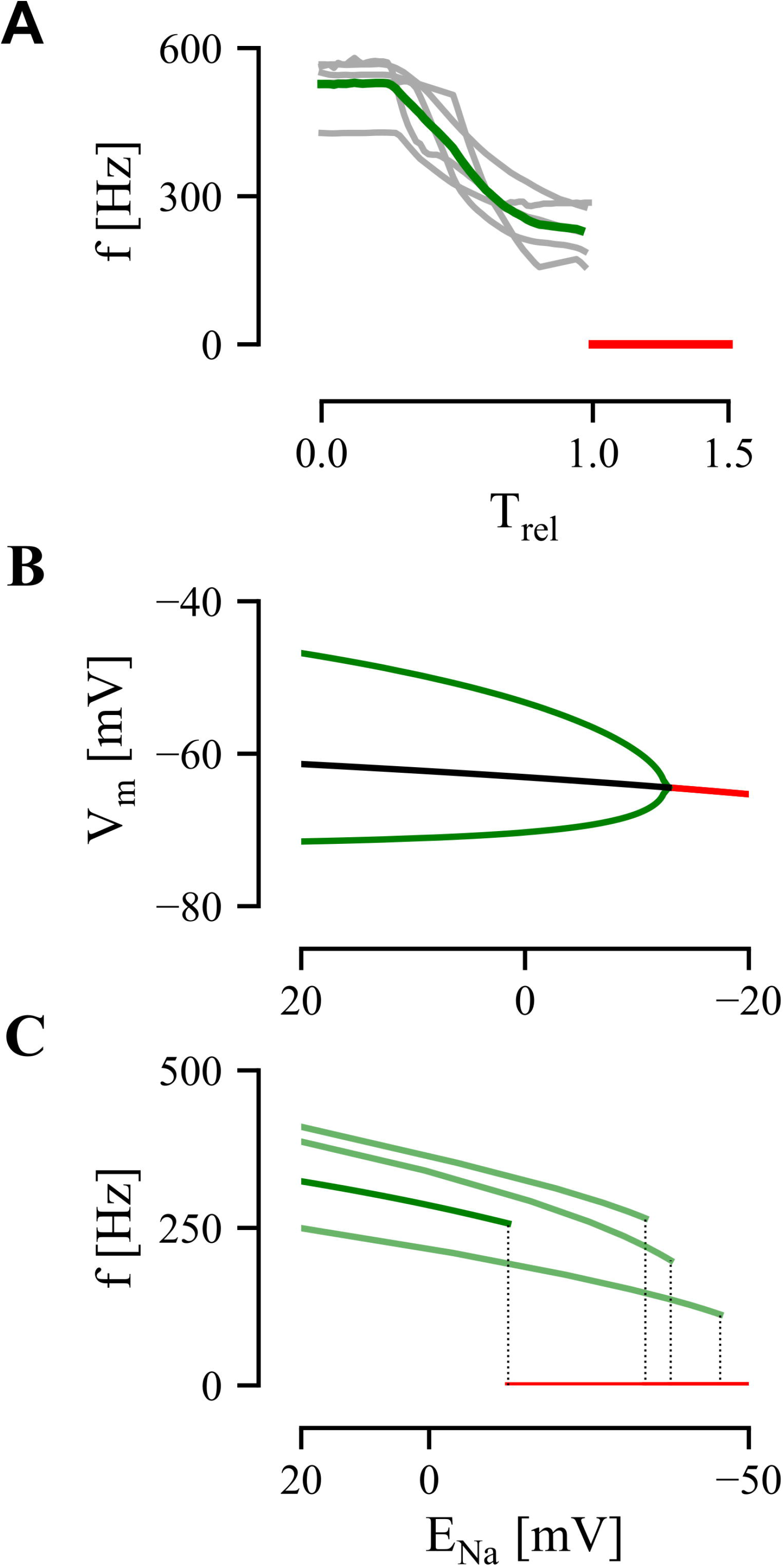
Data and model bifurcation analysis. (A) Time-series of pacemaker frequency (normalized time, T_rel_) as Na-free ACSF is washed in (see Methods) for 5 different pacemaker preparations. Green trace represents average (individual preparations in grey) and red trace represents cessation of firing. T_rel_=1 represents the bifurcation point. (B) Orbit diagram for model bifurcation analysis with respect to E_Na_. Green trace denotes action potential extrema; black trace denotes unstable fixed points; and red trace denotes stable fixed points. Black-Red intersection point denotes the Hopf bifurcation. (C) Frequency analysis of the model Hopf bifurcation. Dark green line shows action potential frequency of the model in figure 1C(i). Light green lines shows firing frequency for other model fits, figure 1C(ii-iv). Red line represents cessation of firing and dotted lines show the bifurcation point of each model.

In figure 3, we show the transition between oscillation and rest in both model and experiments. Our experimental analysis reveals that pacemaker frequency decreases over time, with an abrupt shift to the rest state (for simplicity we define rest to have zero frequency), as extracellular sodium concentration (i.e. E_Na_) decreases (Figure 3A). To account for variability between experiments, we normalize the time scale such that the PN ceases to oscillate at time t=1. Measurements from individual preparations are shown in grey, with the mean shown in green (N=5 fish). On average, we see that the oscillation stops at ~260 Hz (figure 3B).

For the pacemaker model, we compute a bifurcation diagram showing the system’s state (membrane potential, V_m_) for different values of E_Na_ using XPP [34] (figure 3B). As E_Na_ decreases, the model neuron transitions from a oscillating (membrane potential extrema in green) to rest (red) at a bifurcation point corresponding to E_Na_ = −12.8 mV with no hysteresis. To distinguish a Hopf from a homoclinic bifurcation, we measured the frequency of the oscillation as E_Na_ decreases in figure 3C (dark green trace). The oscillation frequency follows the square-root-like curve, characteristic of a Hopf bifurcation [32] until a discontinuity at E_Na_=−12.8mV, after which the cell stops firing. In other words, we can say that as E_Na_ is increased, our model undergoes a supercritical Hopf bifurcation at E_Na_=−12.8mV. We observe qualitatively similar dynamics for the other model fits (figure 1C, light green traces).

While both the models and recordings undergo a sudden loss of spiking, a feature consistent with a Hopf bifurcation, other features of the model dynamics do not match the data. First, the manner in which frequency decreases with decreasing Na^+^ is very different, as indicated by an increasing versus decreasing second derivative (compare figure 3A and 3C). This is possibly due to nonlinear changes in E_Na_ in the experiments resulting from variations in mixing and diffusion through the tissue; this would affect the detailed time-course of E_Na_ but the relation between E_Na_ and time will still be monotone. Secondly, we note that the experimental data spans a larger frequency range (~525Hz→~230Hz; a drop of 60%) whereas the model spans ~380Hz→260Hz, a drop of ~30% (although for the other model fits we see 40-50% drops). This could be due to several factors, but one interesting possibility is that the pacemaker sensitivity to Na^+^ is higher due to the presence of a persistent sodium current that underlies frequency control. It is important to note however, that despite these quantitative differences, both model and experimental systems show similar dynamics, with each exhibiting a Hopf bifurcation.

### Model validation: Pharmacological Manipulations

In a similar manner, we can also consider how PN dynamics change as individual currents are manipulated. Smith and Zakon [27] showed that blocking either I_Na_ or I_K_ channels results in firing cessation in experimental preparations. Importantly, they measured action potential waveform parameters as the channel blocker was washed in (i.e. as an increasing fraction of the channels are blocked). These data can thus provide another means of model validation.

We simulated this gradual channel block in the pacemaker model by manipulating the conductance of the appropriate channel as G_ion_ → (1 − b)G_ion_ where *b* is a number between 0 and 1; control (b = 0) represents no block, while b =1 represents complete block. We demonstrate the effects of progressive block of either Na^+^ (left) and K^+^ (right) channels in figure 4A (block level, b = 0 to b = 0.7). As with the E_Na_ manipulations, we performed a bifurcation analysis in XPP [34] and found that progressive block of both Na^+^ and K^+^ channels is associated with a supercritical Hopf bifurcation (not shown). The reader familiar with bifurcation analyses may note that the decreases in g_Na_ appear to result in a continuous frequency drop, but over a larger parameter range there is in fact a discontinuous drop in frequency associated with a Hopf bifurcation.

**Fig 4.**
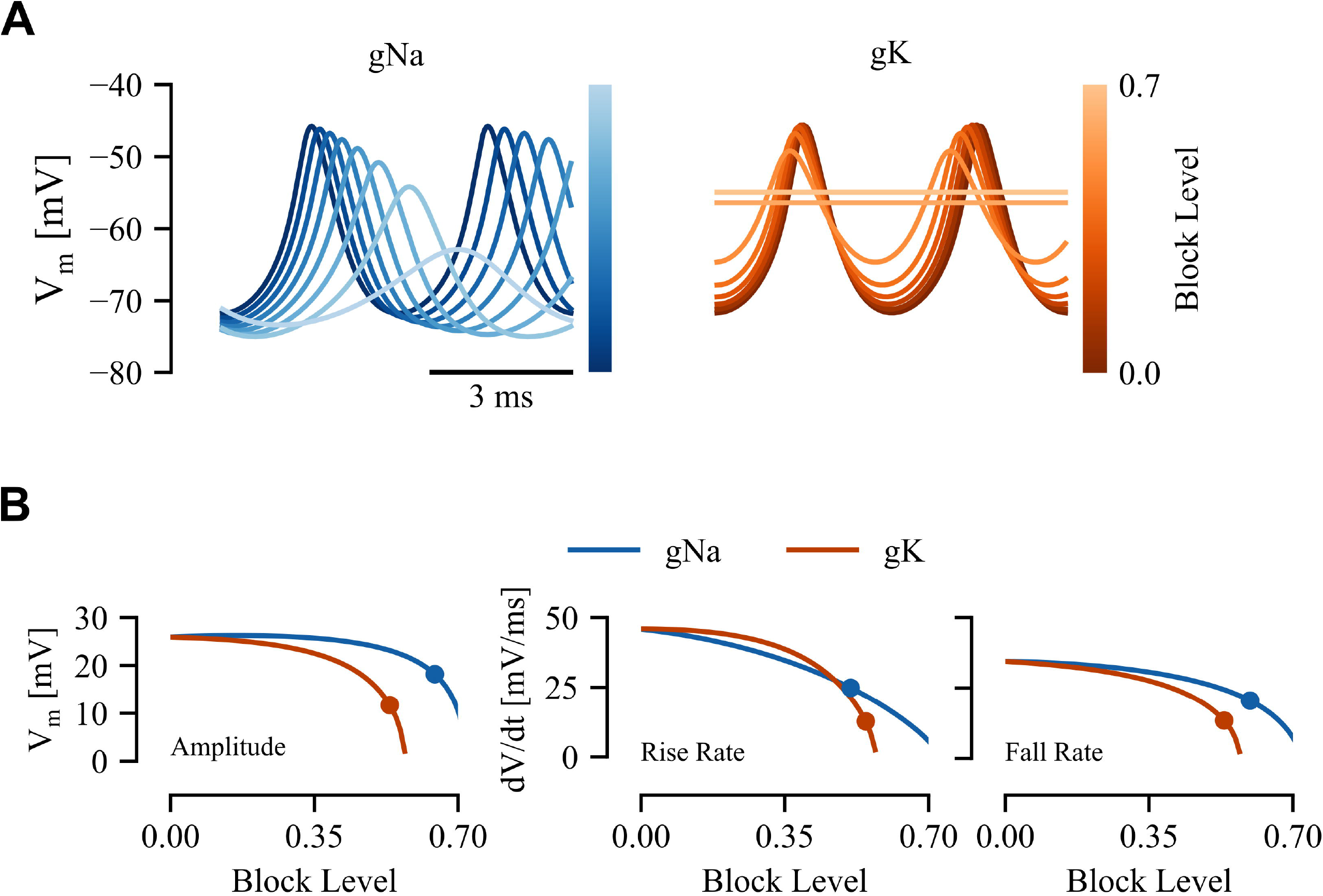
Response to progressive block of Na^+^ and K^+^ channels. (A) Model response to Na^+^ channel block (g_Na_, left) and K^+^ channel block (g_K_, right). (B) Action potential properties measured as a function of block level for peak-peak amplitude (left), action potential rise rate (center) and fall rate (absolute value; right). Dots represent block level with equivalent percentage change in each property from data reported in [27]

Because we do not know the equivalent block level at the time the waveform properties were measured in the experiments, we use the following qualitative comparison based on three action potential waveform features. We consider how these features vary as the block level is increased (figure 4B), and what block level is required to match experimental data (solid circle, figure 4B). A good model will be internally consistent such that the required block level should be consistent across all waveform features. In figure 4B we show the peak-to-peak amplitude, peak rise rate, and peak fall rate (taken as positive for symmetry with rise rate) as a function of Na^+^ and K^+^ channel block level. The equivalent block level (that which corresponds to the percent change noted in the original data) is indicated by a solid circle: the block level for Na^+^ is 0.59 ± 0.06 (mean ± standard deviation) whereas that for K^+^ is 0.55 ± 0.007; the similarity in these values suggests good model performance.

Overall, our analyses show that this new model captures the main oscillatory dynamics and action potential waveforms of pacemaker cells based on the underlying sodium and potassium currents. Further, the results suggest that oscillatory dynamics in pacemaker neurons arise in a specific manner (supercritical Hopf bifurcation) that may influence network synchrony and stability [35, 36].

## Discussion

The pacemaker network (PN) of wave-type electric fish sets the timing of a neural oscillation which exhibits precision and stability far beyond that of any known biological oscillator [13,14]. To understand these dynamics, we have developed a biophysically relevant model of pacemaker neurons that reproduces the action potential waveform as well as the effects of various experimental manipulations. From a dynamical systems perspective, we show that our model undergoes a Hopf bifurcation as E_Na_ is decreased. A similar effect is seen experimentally when Na+ is removed from the extracellular medium: oscillations stop with a minimum frequency ~260 Hz. In these experiments however, we were not able to successfully recover a normal oscillation after low-Na^+^ treatment, and thus could not differentiate between sub and supercritical bifurcations based on the presence of hysteresis. This could be due to a network bistability where the control Na+level could permit oscillation or not. Further work is of course required. We were nonetheless able to show a sudden stop in oscillation which rules-in a Hopf bifurcation and rules out any form of homoclinic bifurcation, where the birth of an oscillation can have an arbitrarily low frequency. In addition, our model accurately reproduces the changes in waveform properties such as amplitude and peak slew rate caused by partial channel block.

Of particular interest in this study is the fact that our model was fit to data from two related species (*A. leptorhynchus* and *A. albifrons*). While there are known differences in cell counts, and frequencies [18,37–39], little is known about the differences in pacemaker network dynamics between these species [14]. We show preliminary data to suggest that both species have similar action potential waveforms (figure 1B) despite wide variations in baseline potential, peak-peak amplitude, and frequency. The implication of this being that pacemaker cells in both species have similar dynamics. This is supported by the fact that the model was fit to these different waveforms within relatively narrow parameter bounds. Furthermore, experimental validations are done with data from both *A. albifrons* (E_Na_) and *A. leptorhynchus* (channel block).

This model can also provide important insight into precision and synchrony in the PN. For example, fast, early currents such as I_Na_p__ tend to decrease synchrony across a network [35] whereas some of the slow, late potassium currents [35, 36]) tend to increase network synchrony [35]. Interestingly, we found that I_Na_p__ was not required to explain pacemaker waveforms, but a delayed potassium current played a fundamental role. Further, the specific dynamics of individual neurons in a network can also influence network synchronization [35, 36]. In particular, neurons exhibiting supercritical Hopf bifurcations (also referred to as Type II excitability) can lead to more robust synchronization [25, 26]. Understanding the role of bifurcation structure in PN precision and synchrony will require future modeling and experimental work.

We acknowledge that our model is a single cell model fit to data from an intact pacemaker network. The impact of this is not clear. In principle, gap junctional strength is proportional to the voltage difference between cells, so in a synchronized network, the voltage difference between cells is low, hence coupling is minimal, with gap junctions serving primarily as an error-correcting entraining force. This suggests that the network effects of gap junctions may be minimal. Nonetheless, given that the model exhibits similar dynamics to those observed experimentally, it will provide a basis for future work focused on how intrinsic neuronal dynamics interact with gap junctional coupling to produce high temporal precision and synchrony in the pacemaker network.

## Author Contributions

ARS and JEL conceived the project. ARS designed the model and performed the simulations with input from JEL. CMB and YS performed experimental recordings. ARS wrote the first draft; ARS and JEL edited the manuscript. All authors have read and approved the final manuscript.

## Supporting information

**S1 Fig. Voltage-dependence functions for each model current across all model fits.** Thick lines represent canonical model in figure 1C(i) and dashed lines represent models in figure 1C(ii-iv). Gating functions (left) and time constant functions (right) with rows representing functions for sodium, potassium, and calcium descending.

**S1 File. Model Equations** Full model equations for pacemaker neuron.

**S1 Table Genetic Algorithm Bounds.** Upper and lower bounds for parameter selection in the GA

**S2 Table Ionic Parameters for model fits.** Canonical Model maps to Figure 1A, Model B-D maps to Figure 1B-D

**S3 Table Gating Parameters for model fits.** Canonical Model maps to Figure 1A, Model B-D maps to Figure 1B-D

## Acknowledgments

This work was supported by an Alexander Graham Bell Canada Graduate Scholarship (CGS-D) to AS from the National Sciences and Engineering Research Council of Canada (NSERC) and an NSERC Discovery Grant to JL (05872).

