## Supplementary material for "Dynamics of a neuronal pacemaker in the weakly electric fish *Apteronotus*": S1 File

$$\frac{dv}{dt} = -I_{Leak} - I_{Ca} - I_{Na} - I_K \tag{S1}$$

$$\frac{db}{dt} = \frac{b_\infty - b}{\tau_b} \tag{S2}$$

$$\frac{dg}{dt} = \frac{g_\infty - g}{\tau_g} \tag{S3}$$

$$\frac{dh}{dt} = \frac{h_\infty - h}{\tau_h} \tag{S4}$$

$$\frac{dm}{dt} = \frac{m_\infty - m}{\tau_m} \tag{S5}$$

$$\frac{dn}{dt} = \frac{n_\infty - n}{\tau_n} \tag{S6}$$

$$\frac{dq}{dt} = \frac{q_\infty - q}{\tau_q} \tag{S7}$$

$$I_{Leak} = G_{Leak}(v - E_{Leak}) \tag{S8}$$

$$I_{Ca} = G_{Ca} b^2 g^2 (v - E_{Ca}) \tag{S9}$$

$$I_{Na} = G_{Na} hm (v - E_{Na}) \tag{S10}$$

$$I_K = G_K n^2 q^2 (v - E_K) \tag{S11}$$

$$\tau_b = \frac{s_{\tau_b}}{\exp\left(\frac{\theta_{\tau_b} - v}{\sigma^2_{\tau_b}}\right) + \exp\left(-\frac{\theta_{\tau_b} - v}{\sigma^1_{\tau_b}}\right)} \tag{S12}$$

$$\tau_g = \frac{s_{\tau_g}}{\exp\left(\frac{\theta_{\tau_g} - v}{\sigma^2_{\tau_g}}\right) + \exp\left(-\frac{\theta_{\tau_g} - v}{\sigma^1_{\tau_g}}\right)} \tag{S13}$$

$$\tau_h = \frac{s_{\tau_h}}{\exp\left(\frac{\theta_{\tau_h} - v}{\sigma^2_{\tau_h}}\right) + \exp\left(-\frac{\theta_{\tau_h} - v}{\sigma^1_{\tau_h}}\right)} \tag{S14}$$

$$\tau_m = \frac{s_{\tau_m}}{\exp\left(\frac{\theta_{\tau_m} - v}{\sigma^2_{\tau_m}}\right) + \exp\left(-\frac{\theta_{\tau_m} - v}{\sigma^1_{\tau_m}}\right)} \tag{S15}$$

$$\tau_n = \frac{s_{\tau_n}}{\exp\left(\frac{\theta_{\tau_n} - v}{\sigma^2_{\tau_n}}\right) + \exp\left(-\frac{\theta_{\tau_n} - v}{\sigma^1_{\tau_n}}\right)} \tag{S16}$$

$$\tau_q = \frac{s_{\tau_q}}{\exp\left(\frac{\theta_{\tau_q} - v}{\sigma^2_{\tau_q}}\right) + \exp\left(-\frac{\theta_{\tau_q} - v}{\sigma^1_{\tau_q}}\right)} \tag{S17}$$

$$b_\infty = \frac{1}{\exp\left(\frac{\theta_{b_\infty} - v}{\sigma_{b_\infty}}\right) + 1} \tag{S18}$$

$$g_\infty = \frac{1}{\exp\left(-\frac{\theta_{g_\infty} - v}{\sigma_{g_\infty}}\right) + 1} \tag{S19}$$

$$h_\infty = \frac{1}{\exp\left(-\frac{\theta_{h_\infty} - v}{\sigma_{h_\infty}}\right) + 1} \tag{S20}$$

$$m_\infty = \frac{1}{\exp\left(\frac{\theta_{m_\infty} - v}{\sigma_{m_\infty}}\right) + 1} \tag{S21}$$

$$n_\infty = \frac{1}{\exp\left(\frac{\theta_{n_\infty} - v}{\sigma_{n_\infty}}\right) + 1} \tag{S22}$$

$$q_\infty = \frac{1}{\exp\left(-\frac{\theta_{q_\infty} - v}{\sigma_{q_\infty}}\right) + 1} \tag{S23}$$
