## Supplementary material for "Dynamics of a neuronal pacemaker in the weakly electric fish *Apteronotus*": S1 Table

| Parameter | Lower Bound | Upper Bound | Unit |
|---|---|---|---|
| $E_{Ca}$ | 20. | 30. | mV |
| $E_K$ | -90. | -80. | mV |
| $E_{Leak}$ | -90. | -80. | mV |
| $E_{Na}$ | 20. | 30. | mV |
| $G_{Ca}$ | 0. | 20. | mS |
| $G_K$ | 30. | 70. | mS |
| $G_{Leak}$ | 0. | 3. | mS |
| $G_{Na}$ | 30. | 70. | mS |
| $s_{\tau_b}$ | 100. | 2. | ms |
| $s_{\tau_g}$ | 5. | 15. | ms |
| $s_{\tau_h}$ | 5. | 15. | ms |
| $s_{\tau_m}$ | 100. | 2. | ms |
| $s_{\tau_n}$ | 5. | 15. | ms |
| $s_{\tau_q}$ | 100. | 2. | ms |
| $\sigma^1_{\tau_b}$ | 10. | 20. | mV |
| $\sigma^1_{\tau_g}$ | 10. | 20. | mV |
| $\sigma^1_{\tau_h}$ | 5. | 15. | mV |
| $\sigma^1_{\tau_m}$ | 5. | 15. | mV |
| $\sigma^1_{\tau_n}$ | 5. | 15. | mV |
| $\sigma^1_{\tau_q}$ | 10. | 20. | mV |
| $\sigma^2_{\tau_b}$ | 10. | 20. | mV |
| $\sigma^2_{\tau_g}$ | 10. | 20. | mV |
| $\sigma^2_{\tau_h}$ | 5. | 15. | mV |
| $\sigma^2_{\tau_m}$ | 5. | 15. | mV |
| $\sigma^2_{\tau_n}$ | 25. | 35. | mV |
| $\sigma^2_{\tau_q}$ | 20. | 30. | mV |
| $\sigma_{b_\infty}$ | 10. | 20. | mV |
| $\sigma_{g_\infty}$ | 10. | 20. | mV |
| $\sigma_{h_\infty}$ | 5. | 10. | mV |
| $\sigma_{m_\infty}$ | 5. | 10. | mV |
| $\sigma_{n_\infty}$ | 10. | 20. | mV |
| $\sigma_{q_\infty}$ | 5. | 15. | mV |
| $\theta_{b_\infty}$ | -70. | -60. | mV |
| $\theta_{g_\infty}$ | -110. | -100. | mV |
| $\theta_{h_\infty}$ | -90. | -70. | mV |
| $\theta_{m_\infty}$ | -70. | -50. | mV |
| $\theta_{n_\infty}$ | -65. | -45. | mV |
| $\theta_{q_\infty}$ | -55. | -25. | mV |
| $\theta_{\tau_b}$ | -100. | -80. | mV |
| $\theta_{\tau_g}$ | -85. | -75. | mV |
| $\theta_{\tau_h}$ | -90. | -60. | mV |
| $\theta_{\tau_m}$ | -90. | -70. | mV |
| $\theta_{\tau_n}$ | -65. | -45. | mV |
| $\theta_{\tau_q}$ | -55. | -35. | mV |

S1 Table
