## Supplementary material for "Dynamics of a neuronal pacemaker in the weakly electric fish *Apteronotus*": S2 Table

| Parameter | **Canonical Fit** | Model ii Fit | Model iii Fit | Model iv Fit | Unit |
|---|---|---|---|---|---|
| $E_{Ca}$ | 23.95 | 22.13 | 29.01 | 27.02 | mV |
| $E_K$ | -80.87 | -87.12 | -84.49 | -89.02 | mV |
| $E_{Leak}$ | -88.91 | -84.63 | -88.95 | -87.81 | mV |
| $E_{Na}$ | 24.22 | 25.56 | 22.12 | 21.06 | mV |
| $G_{Ca}$ | 14.28 | 4.13 | 1.99 | 2.57 | mS |
| $G_K$ | 59.27 | 50.16 | 39.90 | 33.16 | mS |
| $G_{Leak}$ | 1.13 | 1.98 | 1.11 | 2.17 | mS |
| $G_{Na}$ | 63.13 | 52.48 | 48.66 | 61.82 | mS |

S2 Table
