## Supplementary material for "Dynamics of a neuronal pacemaker in the weakly electric fish *Apteronotus*": S3 Table

| Parameter | **Canonical Fit** | Model ii Fit | Model iii Fit | Model iv Fit | Unit |
|---|---|---|---|---|---|
| $s_{\tau_b}$ | 0.62 | 1.38 | 1.65 | 1.07 | ms |
| $s_{\tau_g}$ | 8.28 | 11.36 | 11.95 | 14.02 | ms |
| $s_{\tau_h}$ | 10.29 | 11.36 | 9.71 | 9.62 | ms |
| $s_{\tau_m}$ | 0.50 | 0.47 | 1.08 | 1.33 | ms |
| $s_{\tau_n}$ | 6.56 | 9.69 | 7.18 | 6.35 | ms |
| $s_{\tau_q}$ | 1.01 | 0.72 | 1.15 | 0.96 | ms |
| $\sigma^1_{\tau_b}$ | 11.27 | 11.31 | 13.50 | 18.50 | mV |
| $\sigma^1_{\tau_g}$ | 17.94 | 17.33 | 17.63 | 17.60 | mV |
| $\sigma^1_{\tau_h}$ | 11.15 | 7.27 | 13.49 | 13.01 | mV |
| $\sigma^1_{\tau_m}$ | 11.98 | 7.20 | 8.86 | 8.94 | mV |
| $\sigma^1_{\tau_n}$ | 7.17 | 12.68 | 10.72 | 13.23 | mV |
| $\sigma^1_{\tau_q}$ | 13.14 | 13.41 | 17.87 | 17.79 | mV |
| $\sigma^2_{\tau_b}$ | 12.62 | 15.89 | 17.79 | 18.41 | mV |
| $\sigma^2_{\tau_g}$ | 14.99 | 17.95 | 15.38 | 17.56 | mV |
| $\sigma^2_{\tau_h}$ | 10.26 | 7.80 | 11.14 | 8.17 | mV |
| $\sigma^2_{\tau_m}$ | 13.52 | 7.70 | 12.87 | 14.10 | mV |
| $\sigma^2_{\tau_n}$ | 26.62 | 32.07 | 33.81 | 31.13 | mV |
| $\sigma^2_{\tau_q}$ | 25.15 | 25.97 | 28.51 | 22.07 | mV |
| $\sigma_{b_\infty}$ | 11.55 | 15.12 | 16.80 | 12.37 | mV |
| $\sigma_{g_\infty}$ | 18.38 | 12.71 | 16.72 | 18.55 | mV |
| $\sigma_{h_\infty}$ | 9.48 | 9.03 | 8.51 | 6.92 | mV |
| $\sigma_{m_\infty}$ | 8.78 | 6.91 | 6.33 | 9.08 | mV |
| $\sigma_{n_\infty}$ | 12.05 | 12.99 | 11.33 | 18.22 | mV |
| $\sigma_{q_\infty}$ | 8.03 | 6.71 | 11.40 | 10.39 | mV |
| $\theta_{b_\infty}$ | -67.10 | -64.67 | -67.86 | -65.61 | mV |
| $\theta_{g_\infty}$ | -106.52 | -106.48 | -102.24 | -106.40 | mV |
| $\theta_{h_\infty}$ | -85.67 | -84.66 | -76.30 | -72.08 | mV |
| $\theta_{m_\infty}$ | -55.85 | -66.36 | -58.86 | -55.27 | mV |
| $\theta_{n_\infty}$ | -52.16 | -59.15 | -56.39 | -59.78 | mV |
| $\theta_{q_\infty}$ | -41.48 | -42.43 | -33.52 | -43.99 | mV |
| $\theta_{\tau_b}$ | -83.44 | -96.35 | -88.60 | -94.56 | mV |
| $\theta_{\tau_g}$ | -82.37 | -83.12 | -77.18 | -82.55 | mV |
| $\theta_{\tau_h}$ | -82.53 | -76.68 | -77.66 | -84.61 | mV |
| $\theta_{\tau_m}$ | -77.87 | -85.17 | -72.28 | -85.84 | mV |
| $\theta_{\tau_n}$ | -52.65 | -59.64 | -47.93 | -49.18 | mV |
| $\theta_{\tau_q}$ | -47.45 | -46.91 | -44.41 | -45.09 | mV |

S3 Table
