## Supplementary figures and images for "Dynamics of a neuronal pacemaker in the weakly electric fish *Apteronotus*"

### S1 Fig

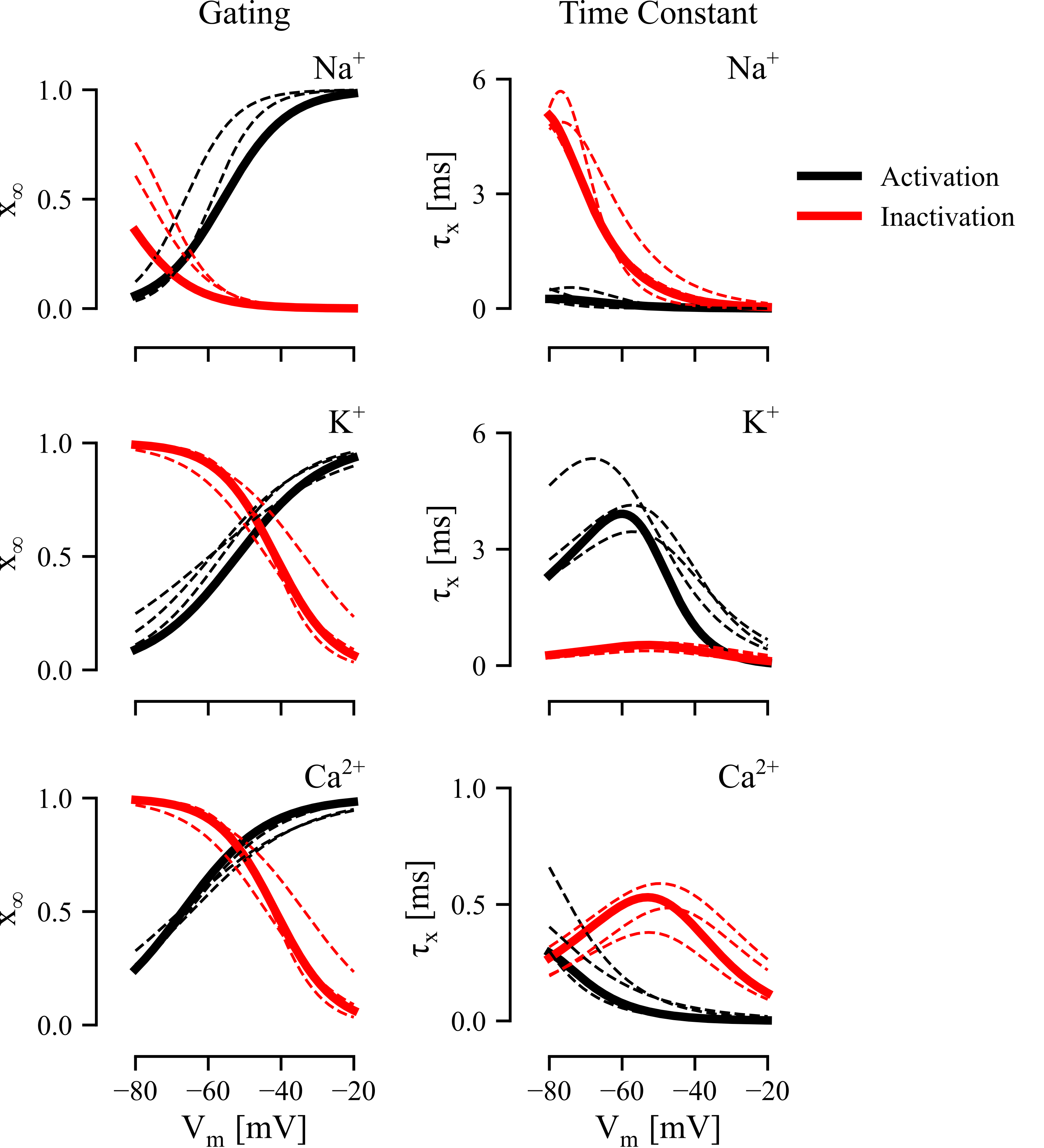
